# MuSCA: a multi-scale model to explore carbon allocation in plants

**DOI:** 10.1101/370189

**Authors:** F. Reyes, B. Pallas, C. Pradal, F. Vaggi, D. Zanotelli, M. Tagliavini, D. Gianelle, E. Costes

## Abstract

**Background and aims:** Carbon allocation in plants is usually represented at a specific spatial scale, peculiar to each model. This makes the results obtained by different models, and the impact of their scale of representation, difficult to compare. In this work we developed a Multi Scale Carbon Allocation model (MuSCA) that can be applied at different, user-defined, topological scales of a plant, and used to assess the impact of each spatial scale on simulated results and computation time.

**Methods:** Model multi-scale consistency and behavior were tested by applications on three realistic apple tree structures. Carbon allocation was computed at five spatial scales, spanning from the metamer (the finest scale, used as a reference) up to 1^st^ order branches, and for different values of a sap friction coefficient. Fruit dry mass increments were compared across spatial scales and with field data.

**Key Results:** The model showed physiological coherence in representing competition for carbon assimilates. Results from intermediate values of the friction parameter best fitted the field data. For these, fruit growth simulated at the metamer scale (considered as a reference) differed from about 1% at growth unit scale up to 35% at first order branch scale. Generally, the coarser the spatial scale the more fruit growth diverged from the reference and the lower the obtained within-tree fruit growth variability. Coherence in the carbon allocated across scales was also differently impacted, depending on the tree structure considered. Decreasing the topological resolution reduced computation time up to four orders of magnitude.

**Conclusions:** MuSCA revealed that the topological scale has a major influence on the simulation of carbon allocation, suggesting that this factor should be carefully evaluated when using different carbon allocation models or comparing their results. Trades-off between computation time and prediction accuracy can be evaluated by changing topological scales.

## INTRODUCTION

Carbon allocation is the process by which the carbon assimilated by leaves, or stored in the form of carbohydrates, is transferred, through the conductive vessels (or phloem tissue), to other plant parts, where is used primarily for respiration and growth. It is generally accepted that the phloem sap moves as a consequence of osmotically generated pressure gradient, as described by the Munch theory (Münch 1930). Gradients are generated by the differences in the concentrations of carbon (C) assimilates between C-sources (mainly leaves, where C-assimilates are loaded into the phloem) and sinks (where C-assimilates are unloaded from the phloem). While the Münch theory has been refined over time (e.g. with the introduction of the leakage-retrieval mechanism, (Thorpe *et al.* 2005) and complementary hypothesis have been proposed to explain the lack of fit with experimental evidences (e.g. sieve tubes decomposed in shorter, overlapping components, at the edge of which solutes are transported at the expense of internal energy) (De Schepper *et al.* 2013), the underlying principles are still considered valid. As such, the C-allocation process is thought to depend primarily on the amount and distribution of available C-supplies and demands along the plant structure, and the possibility of C-supplies to flow via the phloem (Ryan and Asao 2014). In this regard, the case in which C-assimilates can freely move through the phloem, namely that distances have no effect on allocation, is known as the ‘common assimilate pool’ and is mainly considered in small plants such as tomato (Heuvelink 1995) and used in some models (Guo *et al.* 2006; Luquet *et al.* 2006) for grasses.

When plant topology increases in complexity, the modeling approaches to C-allocation present in the literature can be organized in four, partly overlapping, categories (Thornley and Johnson 1990; Lacointe 2000; Le Roux *et al.* 2001; Génard *et al.* 2008): models based on empirical relationships between plant parts and/or the environment; teleonomic models representing the plant as moving towards an a priori defined specific state; source–sink driven: in which different plant components are supposed to attract C-assimilates with different strength; transport resistance and biochemical, representing phloem transport as starting from osmotic gradients and biochemical conversions. The most mechanistic models, representing osmotic flows starting from osmotic gradients, however, imply complex formalizations, highly detailed plant descriptions and because of this received relatively limited attention (Bancal 2002). Conversely, because of their modest mathematical complexity, nonetheless related to a process-based approach, source-sink based models were the focus of a much higher number of studies. In all cases, however, the representation of C-allocation on large tree structures described at high resolution remained computationally unfeasible (Balandier *et al.* 2000; Bancal 2002).

Regarding the spatial representation, each model represents the plant structures, and the process of carbon allocation, at a specific spatial scale. The choice of the scale can be driven by the spatial scope of management or research purposes, the computation time needed for the simulations and the resolution at which the data collected for calibration/model testing were obtained. In general, this spans from the individual metamer (M) (Allen *et al.* 2005) to collections of metamers that, grouped according to criteria such as their age, position or organ type (Kang *et al.* 2008), constitute larger portions of a tree structure, such as branches, main axis and so on, until whole plant compartments (Lakso and Johnson 1990; Kang *et al.* 2008). In this context, it is here worth mentioning a few source-sink based models, for later considerations. The L-PEACH model (Grossman and DeJong 1994; Allen *et al.* 2005) is a reference for completeness of the described processes and represents tree growth over multiple years. This uses a transport resistance analogy to mimic C-transport at the M scale (internode, leaf, fruit). The QualiTree model represents growth and quality of fruits on a peach tree structure, at the fruiting unit (FU) scale, during a single growth season (Lescourret *et al.* 2011). The SIMWAL model represents the growth of a young walnut tree organized in axes, which are divided into growth units (the scale of C-allocation), in turn are split into internodes and nodes (Balandier *et al.* 2000).

The computation of C-allocation by a model is affected by both the formalism used to describe the physiological processes and the discretization of the plant. Because of this, isolating individual effects for model inter-comparison is hardly achievable.

The present work proposes a new, multi-scale carbon allocation model (named MuSCA), whose aim is to allow for a flexible definition (possibly based also on non-topological elements, such as age or organ type) of the topological plant scales (also proxy of spatial resolution), concomitantly with the simulation of carbon allocation, rather than choosing, a priori, one single spatial scale. The model relies on the use of Multi-scale Tree Graph (MTG) (Godin and Caraglio 1998), a formalism inspired by the observation of the multi-scale organization of plant structures (Barthélémy 1991; Barthélémy and Caraglio 2007; Balduzzi *et al.* 2017). MTG allows for the topological description of a plant at multiple, nested, spatial scales on the same graph. In addition to the connections and boundaries of the described topological scales, intensive (e.g. organ type) or extensive (e.g. geometrical features) properties can be referred to the individual plant component at any scale. MTG has been previously used in plant architectural models (Costes *et al.* 2008) and in radiation interception models (Da Silva, Han, and Costes 2014), as well as to simulate physiological processes (Fournier *et al.* 2010) or to model plant-pathosystems (Garin *et al.* 2014, 2018; Robert *et al.* 2018) but, in the latter case, by using one scale at a time. Differently, in MuSCA the topological scale at which C-sources and sinks are computed can change based on user’s choice, prior to individual cycles of carbon allocation. As such, C-allocation can be computed on the same plant structure represented at different topological scales.

In this paper, we first provide an overview of the model, define its scope and inputs, present the formalisms used to compute C-allocation and to move across scales (Model Description). Afterward, the model is calibrated for the apple tree (*Malus* x *domestica Borkh.*), Fuji cultivar, for C-demands, and linked to a radiative model for the estimation of C-assimilates on plants represented as MTGs (Application to realistic tree structures: the case of apple (Malus domestica)). The model is then applied on three contrasted tree structures produced by an architectural model, and for different sap friction parameters, and the emerging effects of competition for C-assimilates on fruit growth are analyzed (Results, Discussion). The fruit growth distributions simulated on one structure for different sap friction parameter values are compared to field observations to retain the parameter value that might best represent sap flow dynamics. The coherence of the multi-scale formalism and the influence of the scale of representation on the simulated carbon allocation are analysed. Finally, the trade-off between computational efficiency and accuracy are discussed, also in respect to the possible interactions between specific tree structure and the topological scale used.

## MATERIALS AND METHODS

### Model Description

#### Overview

MuSCA is a generic, functional structural plant model (FSPM) (White and Hanan 2012) representing the carbon allocation and growth of a plant at different, user-defined, spatial scales, while taking into account competition and distances between carbon sinks and supplies present on the plant (Fig.1). The model is designed as a set of modules. In this modular architecture, each module is generalized so that only a few of them are specific to a given cultivar or species (Table 1, Table 2). The model is integrated in the open-source OpenAlea environment (Pradal *et al.* 2008, 2015) and implemented in the Python language. It makes large use of other components available in OpenAlea, such as the MTG dynamic data-structure (https://github.com/openalea/mtg), used to represents the topological connections in plant structures, the PlantGL library (Pradal *et al.* 2009) used to represent the 3D geometry, and the RATP radiative model (Sinoquet *et al.* 2001). In its current version, the MuSCA model simulates biomass accumulation, but not shoot elongation, during a single vegetative season.

**Fig.1:**
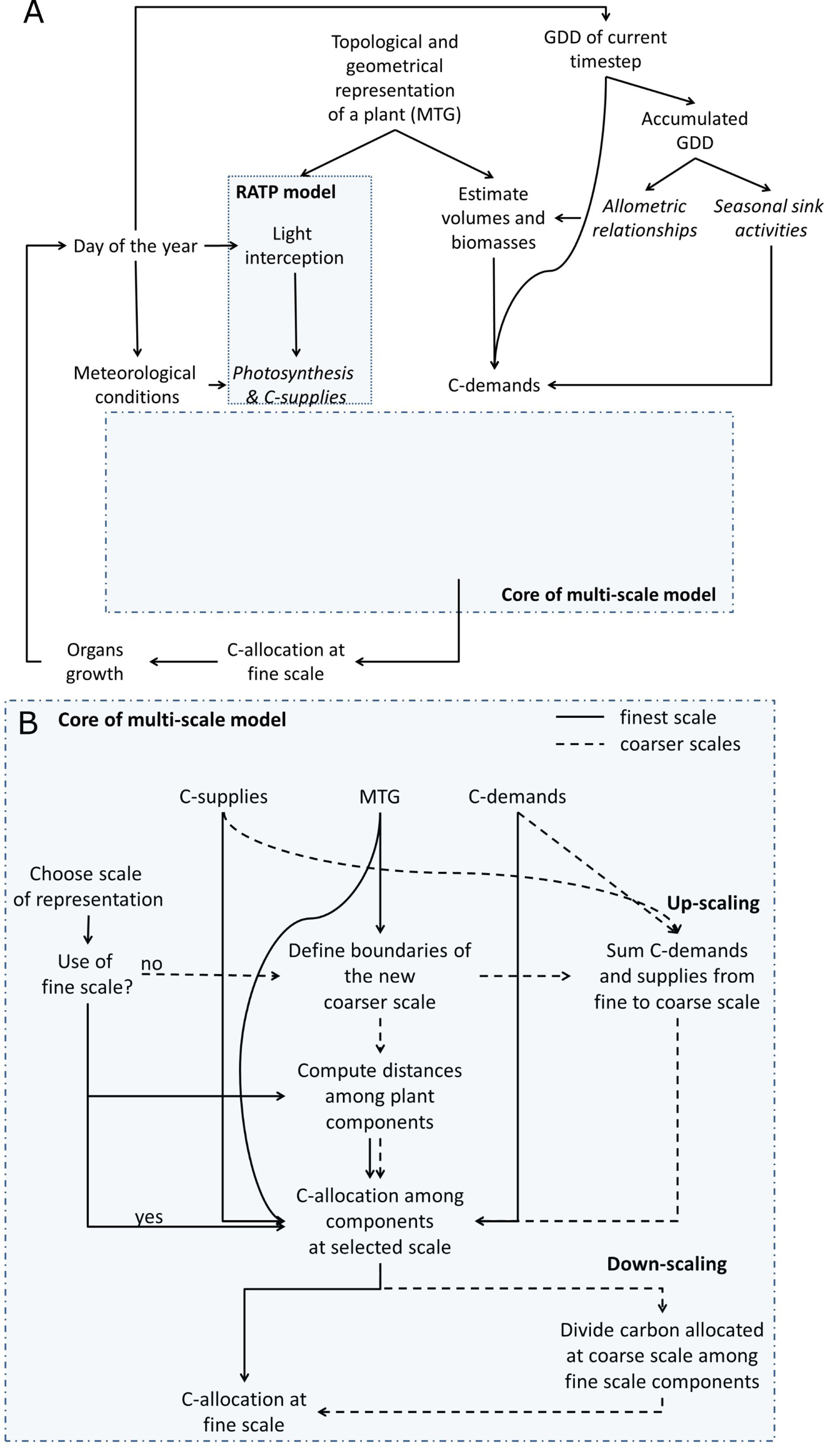
Conceptual workflow of the model: (A) General framework for the application of the model; (B) the core of MuSCA. (A): A multi-scale tree graph representing a plant structure at a given developmental stage is provided. This is used, together with the Growing Degree Days (GDD), allometric relationships and plant geometric information to compute initial volumes and biomasses of plant components at fine scale (metamer). Carbon demands are estimated at metamer scale by making use of species specific seasonal sink activities, biomasses and GDD at the current time step. The position of individual leaves on the plant structure is used by the radiative model RATP to compute light interception and, together with meteorological conditions, the individual leaf photosynthesis. (B): The workflow follows different paths depending on the selected scale: finest scale (solid lines) and coarser scales (dashed lines). The scale at which the plant is represented determines up-and down-scaling processes. In case a coarse scale is chosen, carbon demands and supplies are up-scaled. Carbon allocation is then computed among the plant components at the chosen scale. After allocation, if the selected scale was a coarse one, the carbon allocated to each component is divided among its constituent elements. Biomass of individual elements is eventually updated prior to moving to the next time-step. MTG: Multi-scale Tree Graph. In *italics* are the steps containing species specific parameters.

**Table 1:**
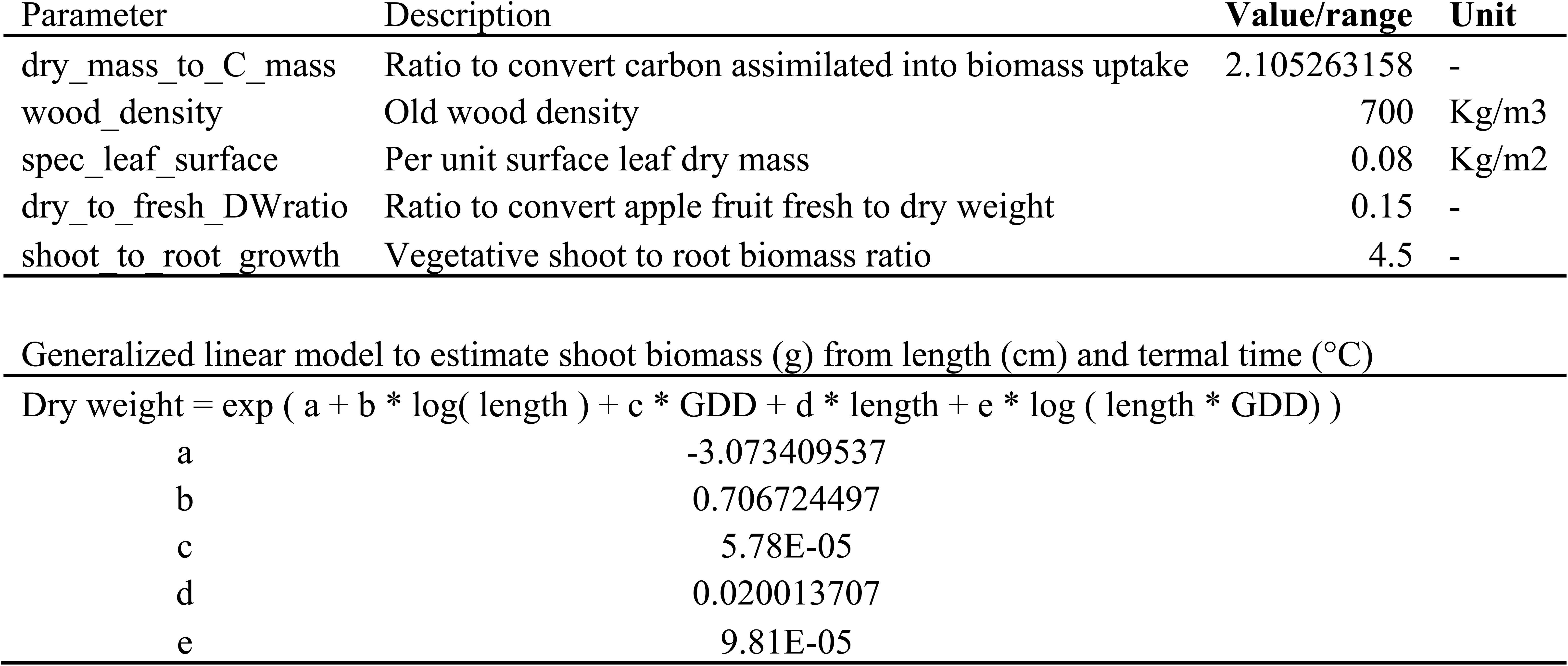
Model parameters and equation for estimation of biomasses.

| Parameter | Description | Value/range | Unit |
| --- | --- | --- | --- |
| dry_mass_to_C_mass | Ratio to convert carbon assimilated into biomass uptake | 2.105263158 | - |
| wood_density | Old wood density | 700 | Kg/m3 |
| spec_leaf_surface | Per unit surface leaf dry mass | 0.08 | Kg/m2 |
| dry_to_fresh_DWratio | Ratio to convert apple fruit fresh to dry weight | 0.15 | - |
| shoot_to_root_growth | Vegetative shoot to root biomass ratio | 4.5 | - |

|  |  |
| --- | --- |
| a | -3.073409537 |
| b | 0.706724497 |
| c | 5.78E-05 |
| d | 0.020013707 |
| e | 9.81E-05 |

**Table 2:**
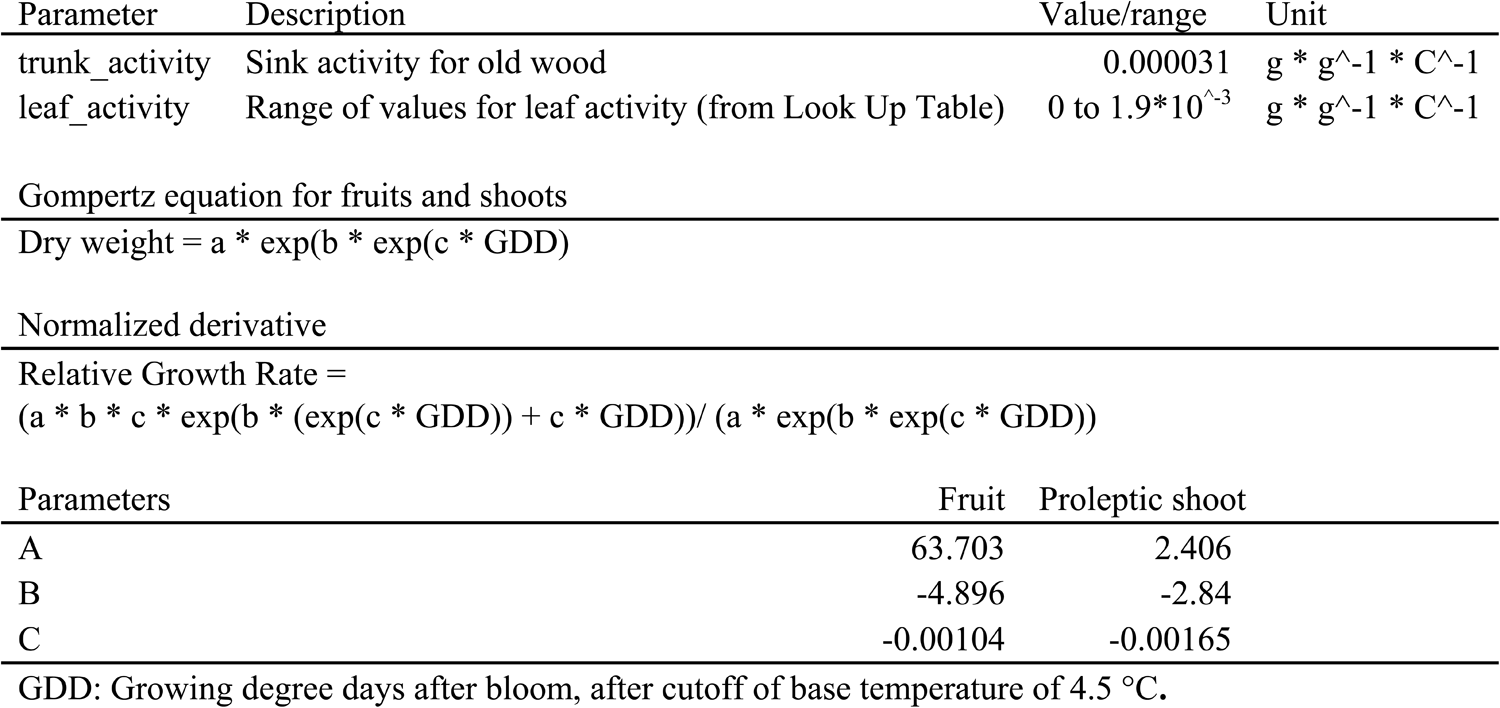
Model parameters and equation for estimation of sink activities

#### Input plant structures and creation of new scales

The input plant structures of MuSCA are MTGs containing a geometric and topological plant description. In this, individual plant components are characterized by qualitative (e.g. the type of connection of the vertex with its parent: branching or succession; the organ type) or quantitative information (geometry or age of plant component). This information can be used to create criteria defining the membership of individual plant component to larger groups of adjacent plant components. In particular, the criteria can be used to just define the edges of the groups, while all the elements in between two edges will be considered as belonging to the same group. These groups correspond to a topological scale coarser than the original one (for an application see Definition of topological scales).

#### Movement of carbon

The movement of the available carbon (*F_ij_*), from a C-supply (*i*) to a C-sink (*j*), is represented as a function of the C-available in the supply (*ACP_i_*), the sink C-demand (*Demand_j_*) (eq. 1a), and as inversely related to the distance (*dist*) and resistance to the flow (*h*, called friction parameter in the following) between components (eq. 1b), along the plant topology. This equation is inspired by a previously defined equation (SIMWAL model, (Balandier *et al.* 2000).

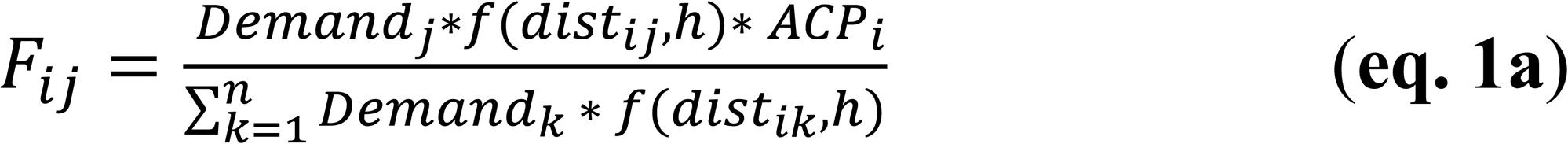

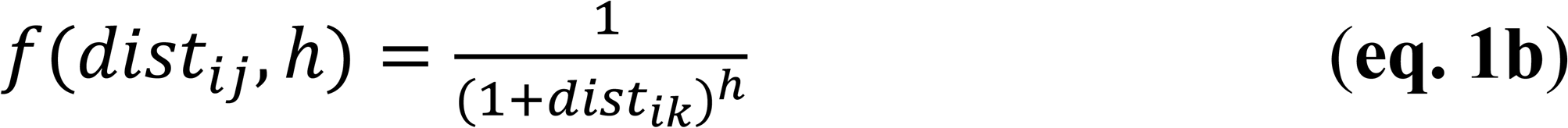

If in excess, the C-allocated to a plant component can be reduced to just fulfill its C-demand, while the excess will be considered as a supply provided by that component on the following time-step (Balandier *et al.* 2000).

The carbon allocated to a topological element is eventually divided among the different organs that constitute it (at the metamer scale: fruit, leaf, internode), proportionally to their individual carbon demand.

Sink C-demand, in turn, are estimated as a function of organ type (fruit, vegetative shoot, leaf, old wood), thermal time expressed as Growing Degree Day (GDD), and organ dry weight (eq.2) (Marcelis 1996), at the daily time-step and at the metamer scale.

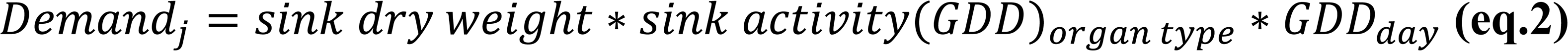

#### Calculation of distances

The use of a source-sink based formulation, where the flow is a function of distance, implies that the distances between individual sources and sinks need to be considered. However, a representation of the C-allocation process coherent at multiple topological scales requires the definition of a multi-scale equation of distances, independent of the scale of representation.

The computation of the C-allocation process is computationally demanding because it requires to compute the distance between all the pairs of carbon sinks and sources in the tree which, in a plant composed of n vertices, is comparable to n**2 elements. Considered this, we propose to define an algorithm, based on the multi-scale organization of plants, that allows reducing the number independent plant vertices, and thus the operations needed to compute the distance between them. In order to do so, we formulate a new equation to compute distances suitable at multiple scales.

The distance (*dist_i,j_*) between a source and a sink vertex (*i*, *j*) of an MTG is defined by the length of the topological path connecting source and sink vertices. As such, the distance is defined as the sum of the euclidean distances (*D*) between the base and the barycenter (semi-length) of each extremity (*i* and *j*), plus the *D* between the successive basis of the plant components (evaluated at the selected scale) connecting (in topological sense) the two vertices (eq. 3a, 3b, 3c, Fig.2). In order to evaluate these distances: first, the greatest common ancestor (*GCA*) between the two vertices is identified. This is done by taking each of the two vertices and considering their parents iteratively (its parent, the parent of the parent,…). The *GCA* is identified as the first parent being ancestor of both vertices (Fig.2). In case the considered vertex is the base of the mtg, this is directly considered as the *GCA*. Second, the *D* occurring between the bases of successive vertices, along the paths connecting the input vertices to their *GCA* (excluded), are summed up (eq. 3d). In addition, if the *GCA* is not any of the two input vertices, the *D* between the bases of the two vertices (insertion points in the *GCA*) is also summed (eq. 3e).

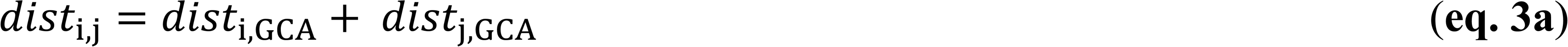

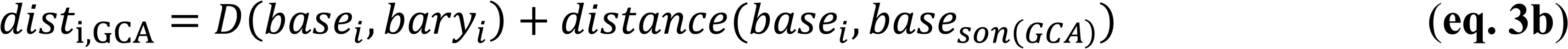

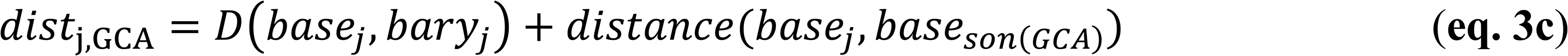

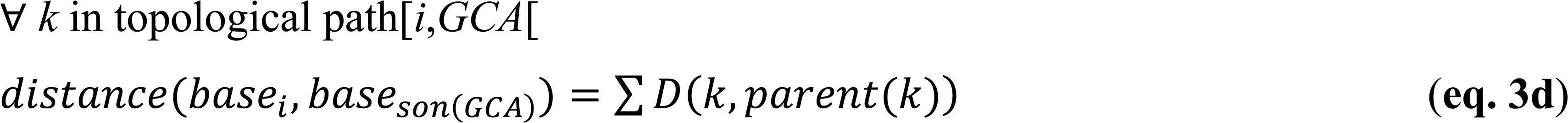

if i,j ⊆ successors(*GCA*):
for *son(GCA)i* ⊆ in topological path [*i*, *GCA*] and son(GCA)j ⊆ topological path [*j*, *GCA*]

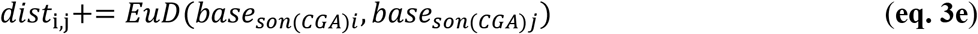

**Fig.2:**
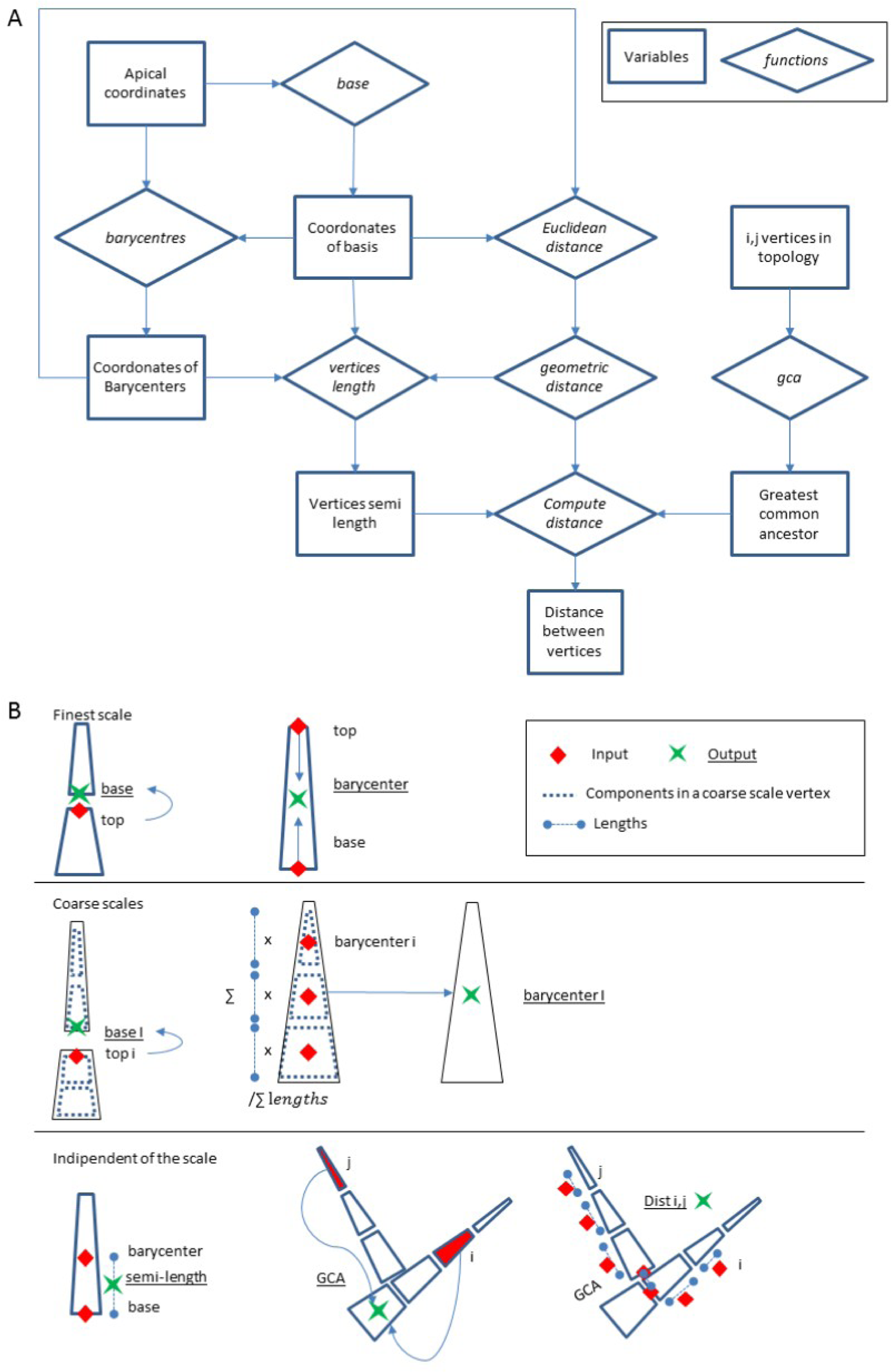
Workflow and graphical representation of the calculation of distances. (A) Workflow for the calculation of distances. (B) Graphical representation of the: (above) calculation of coordinates of basis and barycenters at finest and (middle) coarser scales. (Below) calculation of the semi-length of components, identification of greatest common ancestors and computation of distances along a path connecting two vertices. At coarse scale (middle), low case letters indicate finest scale, while capital letters refer to coarse scale vertices. GCA: Greater Common Ancestor.

The spatial coordinates of basis and the barycenter of individual vertices are found differently depending on the selected scale: regarding the vertices at the finest scale, basal coordinates are the top coordinates of the parent vertex (eq. 4a). An exception is the base of the mtg, for which basal coordinates are stored in the complex of the vertex itself (the only vertex not having a parent). Coordinates of barycenters at the finest scale are computed as the mean between the top coordinates of consecutive vertices (example for x-coordinates, eq. 4b) or, for the base of the mtg, as the mean of its top and basal coordinates. Regarding vertices at a coarse scale, as mentioned above, the basal coordinates of the MTG are already available in its base at coarse scale. For any other coarse scale vertex (*I*), given its components (*i*), the one whose parent is not itself a component of the same coarse scale vertex is identified (the basal component of I) (Fig.2B). The top coordinates of its parent are the basal coordinates of the coarse scale vertex (eq. 4c). The coordinates of the barycenter of coarse scale vertex are, instead, calculated as the mean of the coordinates of the barycenters of its components, weighted by their individual lengths (as in eq. 4d for the x coordinate).

At finest scale:

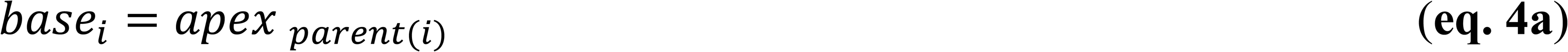

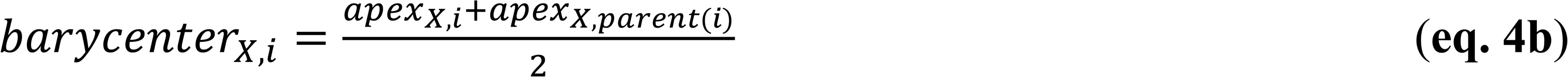

At coarse scales:

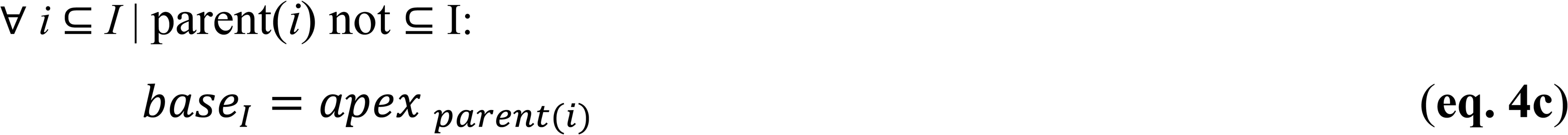

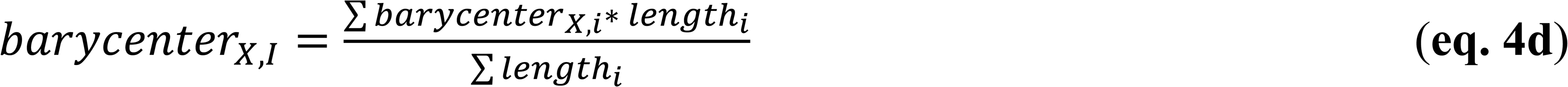

#### Up-and down-scaling

In MuSCA, some of the plant properties available at one scale are used to provide a description of the same properties at a coarser (up-scaling) or finer (down-scaling) scale. This is done in simulations running at coarse scales, prior and after the calculation of carbon flows, respectively by calling up-and down-scaling functions (except for the initial calculation of biomasses, see par From geometry to biomass). The up-scaling of properties such as biomass, carbon supplies and demands of plant parts is simply done by summing up the property values stored in all the vertices that are contained within the topological boundaries of the coarser scale component. Conversely, down-scaling the same properties from a coarse scale component to its constituting elements requires some assumptions. In particular, when the carbon allocated to a coarse scale component is down-scaled, we opted for two options: it can be either assigned proportionally to the relative carbon demands of its constituent elements (default), either equally assigned among them.

##### Application to realistic tree structures: the case of apple (Malus domestica)

In order to assess the model coherence at multiple scales, its physiological soundness and the effect of the scale on computation time, we decided to apply the model on different tree structures of a major fruit tree crop, the apple tree, represented at multiple topological scales (Input tree structures).

For this reason, we developed a few modules for the application of the carbon allocation model on tree structures. Despite their genericity, these modules had to contain some species specific parameters and allometric relationships (Table 1, Table 2). First we developed a module to estimate biomasses of an input tree structure based on its geometrical description and allometric relationships (From geometry to biomass). Second, the C-supplies and demand of the tree are estimated (Source and Sink strengths). Third we defined five topological scales of representation of the tree (Definition of topological scales). Finally simulations of C-allocation and fruit growth were run (Simulation of fruit growth).

#### Input tree structures

In the current application, input tree structures were MTGs produced by the MAppleT model (Costes *et al.* 2008). This model includes Markov models for simulating annual shoot branching and successive growth across years, a biomechanical model simulating the change in branch form over time, and a modified pipe-model to estimate axis radial growth. The leaf area and internode length depend on their rank along the shoot (Da Silva, Han, and Costes 2014). The MTGs output produced by MappleT are apple trees structures at a certain stage of development (given date of the year), represented at two scales, the growth unit and metamer, as well as the 3D coordinates of each metamer and leaf area for the leafy shoots.

We used one simulated apple tree (‘Fuji’ cultivar) that was four years old, and two-three-years old other apple trees, that were previously generated for a sensitivity analysis (Da Silva, Han, Faivre, *et al.* 2014). These two apple trees (named afterwards: Ap-05 and Ap-10), were chosen as well branched trees, having different vigor and size. The three simulated trees represented tree structures in late June (day of the year = 182), namely after the end of shoot elongation. On them, initial individual fruit dry weight was set identical for all fruits and equal to 8 g (Reyes *et al.* 2016).

#### From geometry to biomass

At the beginning of the simulation, the model uses the geometrical description of the plant and some species specific parameters to estimate the initial dry weight of each plant component at the finest scale (in the current application M scale: internode possibly with a leaf and/or a fruit) (see Supplementary Information: Inputs of the MuSCA model.). In particular: dry weight of old wood and vegetative internodes are computed as functions of their geometrical description (lengths and radiuses), and of a constant (wood density) or a function of thermal time, respectively; fruits dry weight is calculated from fresh weight and a constant (dry to fresh fruit dry weight ratio); leaf dry weight is calculated from surface and a constant (per unit surface leaf dry mass) (Table 1). The multi-scale feature of MTG is used to calculate the dry mass of internodes starting from the dry mass estimated for the current year shoot to which they belong. In particular, volumes of internodes are first computed as truncated cones, summed up to provide the volume of individual shoots and stored at a “current year shoot” coarse scale. The length of current year shoots is then obtained by summing the lengths of their internodes, and used to estimate shoot biomass by means of a thermal-time dependent allometric relationship (Table 1) (Reyes *et al.* 2016). Finally, the dry biomass of individual internodes is calculated as their individual volumetric fraction in the shoot to which they belong, multiplied by the shoot dry biomass.

Regarding the root, a rough representation of this compartment can be optionally added to the plant. In this case, a root mass proportional to the total mass of the current year shoots is added to the root metamer following the shoot/root functional balance assumption (Davidson 1969; Grechi *et al.* 2007). The length of the roots is represented as equal to half the average distance between the soil and the vegetative shoots. The root basal coordinates are thus defined equal to the tree basal coordinates, except for the vertical (z axis) component, to which the calculated distance is subtracted (downward translation).

#### Source and Sink strengths

In MuSCA, the amount of C-assimilates available from photosynthesis in individual leaves is estimated by means of a link to a radiative model. The RATP: Radiation, Assimilation, Transpiration and Photosynthesis, turbid-medium based, model (Sinoquet *et al.* 2001) was chosen because of its availability on the OpenAlea platform, the possibility of relatively high spatial resolutions and of its application on MTGs (see also Supplementary Information: Inputs of the MuSCA model.). In RATP the tree structure is first discretized into voxels of user-defined size. The voxel specific mean leaf area density (turbid medium assumption) is calculated based on the plant 3D representation in space. This is then used to compute the direct and diffused PAR and NIR light intercepted in each voxel, and its related photosynthesis, every 30 minutes, across the whole day. The C-assimilation estimated per leaf unit surface is first associated to each leaf and integrated over the whole day, and then converted into dry matter uptake per leaf per day. Optionally, for testing purposes, a constant C-assimilation per unit surface can be used instead of the value provided by the radiative model. In this study, a previous calibration of RATP for the apple trees, Fuji cultivar (Massonnet *et al.* 2006), was used for the computation of carbon assimilation per leaf area unit, depending on the climatic conditions.

Sink strength were estimated based on the analysis of maximum potential growth of the different organs of apple trees of the same cultivar (Table 2). Maximum potential growth curves were obtained by fitting thermal-time-dependent gompertz functions to organ-type-specific maximum dry weights (Grossman and DeJong 1995) of proleptic and epicormic shoots, and fruits of apple Fuji trees growing in conditions where competition for carbohydrates was minimized (Reyes *et al.* 2016). Their normalized derivatives (Hunt 1982; Grossman and DeJong 1995) represents the seasonal patterns of organ type specific activities (or maximum potential relative growth rates). Regarding the old wood, the slope of a linear relationship fitted through the logarithm of the relative growth rate of the old wood biomass vs growing degree days (GDD) obtained by Reyes et al. (2016) was used.

#### Definition of topological scales

For demonstrative purposes we present here five biologically relevant scales of representation of trees. Two of them are commonly used in MTG (metamer and growth unit) and three are newly defined (trunk, branches and shoots; 1st order branch and inter-branches; fruiting unit) (Fig.3). The finest scale considered was the metamer (M) (or phytomer) that is composed of a node and its leaf(ves) and axillary bud(s) plus the subtending internode and constitutes the basic element of plant construction (Barlow 1989; Costes *et al.* 2006; White and Hanan 2012). This is also the spatial scale used in the L-Peach model for carbon allocation (Grossman and DeJong 1994) and it was thought as a reference for the representation of carbon allocation in this study. The second scale corresponds to growth unit (GU) which includes adjacent plant metamers that grew without interruption during a vegetative period (Barthélémy 1991). Because of their age and leafy status, the internodes of an annual shoot, after cessation of the primary growth, could be thought as having similar behavior with respect to the use of assimilates. The third scale corresponded to the discretization of the tree in main trunk, first order branches originating from it and the leafy shoots (Trunk, Branches and Shoots, TBS). The fourth scale corresponded to first order branches originating from the main trunk, but without considering the individual shoots separately (BR1). The fifth scale corresponded to the fruiting unit (FU). A FU includes the set of shoots born from a section of one year old wood, followed by a terminal leafy or bourse shoot. In all scales, the root is considered as a compartment in itself. In scales three and four, the whole trunk is considered as a compartment.

**Fig.3:**
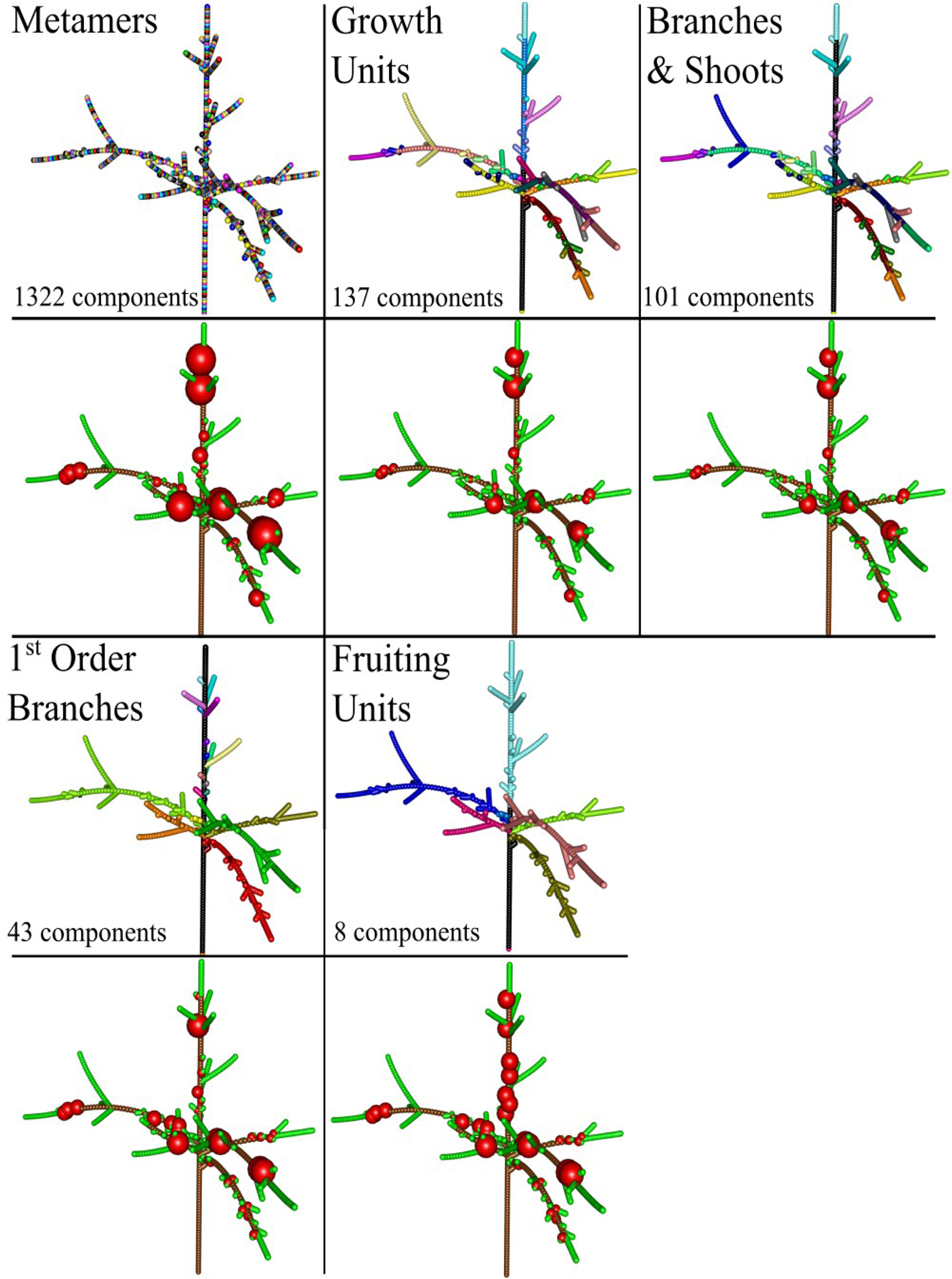
Simulation of carbon allocation and fruit growth on a tree structure represented at different spatial scales during one day. Upper panels: tree parts represented with the same color belong to the same scale, i.e. trees with an higher number of colors are made of a higher number of components. The number of components at each scale is indicated in the corresponding low left corner. Lower panels: the volume of the represented spheres is proportional to the normalized increment in fruit dry weight within the tree. Simulations were run with a friction parameter equal to eight.

#### Simulation of fruit growth

We run simulations for the three apple tree structures represented at the five over-mentioned topological scales. In order to identify a biologically sensible value of the friction parameter (h), this was varied across a range of almost two orders of magnitude (0.5-16).

In a first set of simulations the amount of carbon assimilates was generally higher than the carbon demand at tree scale. This might be due to the fact that the sink activities were possibly underestimated due to the lack of C-loss via respiration. The respiration process is indeed not yet implemented in the in the current version of MuSCA model. As such, in order to analyze the effect of within tree organ competition for C-allocation, simulations were re-run after having artificially doubled all C-demands.

We then tested the model behavior run at M scale by analyzing the growth of individual fruits in relation to both the fruit load and carbon assimilated in surroundings of increasing radiuses, centered on the targeted fruit. The ratio between the amount of C-assimilated and the number of other fruits in the surrounding of individual fruits was calculated for neighborhoods of incremental radii (from 5 cm to 135 cm). This ratio (C-assimilated / number of fruits) represents a comfort index with respect to the availability of C for the fruit and the occurrence of competition with other fruits. The ratio was then correlated to the simulated fruit growth, and its significance adjusted for multiple comparisons with the Bonferroni correction. Biological relevance was also tested by comparing the simulated and harvested fruit size distribution. In particular, based on the assumption that fruit weight obtained in early growth stages is correlated to the fruit weight at harvest (Stanley *et al.* 2000), we compared the distributions of the normalized values of the carbon allocated in our simulations with the one of fruit weight measured at harvest in the field on four-years old Fuji trees (Costes, unpublished data). Similarity among distributions was also assessed in terms of Root Mean Squared Error (RMSE) between distribution counts. Based on these results, a narrower range of friction parameter values was identified as more biologically relevant and used in further analysis.

The coherence of simulations across scales was tested by correlating the fruit growth obtained at each coarse scale with the one at M scale, used as a reference. Results obtained for equal friction parameter and same tree structure were considered together. For each coarse scale component, the average of the fruit growth obtained at M scale within the boundaries of the coarse scale component was computed. As such, a one-to-one comparison was possible between mean growth at M and other (coarser) scales. Deviations between M and the other scales were assessed visually on correlation plots, and by means of the coefficient of variation of the root mean squared error (CV RMSE).

The effect of using different topological scales on the number of plant components and on simulation time was also analyzed. All simulations were run on a personal computer equipped with an Intel i7-6700HQ 2.59 Ghz CPU, 8Gb RAM; OS: Windows 10 Home, 64bit.

## RESULTS

### Testing physiological assumptions

Relative growth rates (RGR) results, aggregated at the compartment level, increased with the friction parameters in the case of shoots while decreased for fruits and old wood (Table 3), as a consequence of the increased tendency of the carbon assimilates to remain closer to the C-source.

**Table 3:** Relative growth rate at compartment scale obtained at metamer scale for different friction parameters (h) on the Fuji tree structure. Figures are reported with five decimals in order to see RGR differences when using different h parameters.

| Relative growth rate (mg g<sup>-1</sup> dd<sup>-1</sup>) |  |  |  |
| --- | --- | --- | --- |
| h | Old wood | Shoot | Fruit |
| 0.25 | 0.0441 | 1.0784 | 2.3105 |
| 0.5 | 0.0447 | 1.0778 | 2.3103 |
| 1 | 0.0456 | 1.0771 | 2.3101 |
| 2 | 0.0465 | 1.0790 | 2.3099 |
| 4 | 0.0443 | 1.1110 | 2.3051 |
| 8 | 0.0371 | 1.2527 | 2.2003 |
| 16 | 0.0269 | 1.2678 | 1.7940 |

Simulation results at M scale show the impact of the friction parameter (h) in determining the area around an individual fruit relevant in terms of competition for C-assimilates with other fruits (Fig.4A,B). For relatively small values of the friction parameter (0.5 - 4), the neighborhood within which the individual fruit growth is significantly correlated to the ratio between C-assimilated and number of fruits is relatively large (>0.9 m). This means that the variability in fruit growth is mostly related to C-sources located far from it. In other terms, a large part of the tree structure affects the growth of each individual fruit. Conversely, for high friction parameter values (8, 16), fruit growth is affected mainly by the C provided by closer leaves and the possible competition by neighboring fruits (neighborhood < 0.9 m) (Fig.4B).

**Fig.4:**
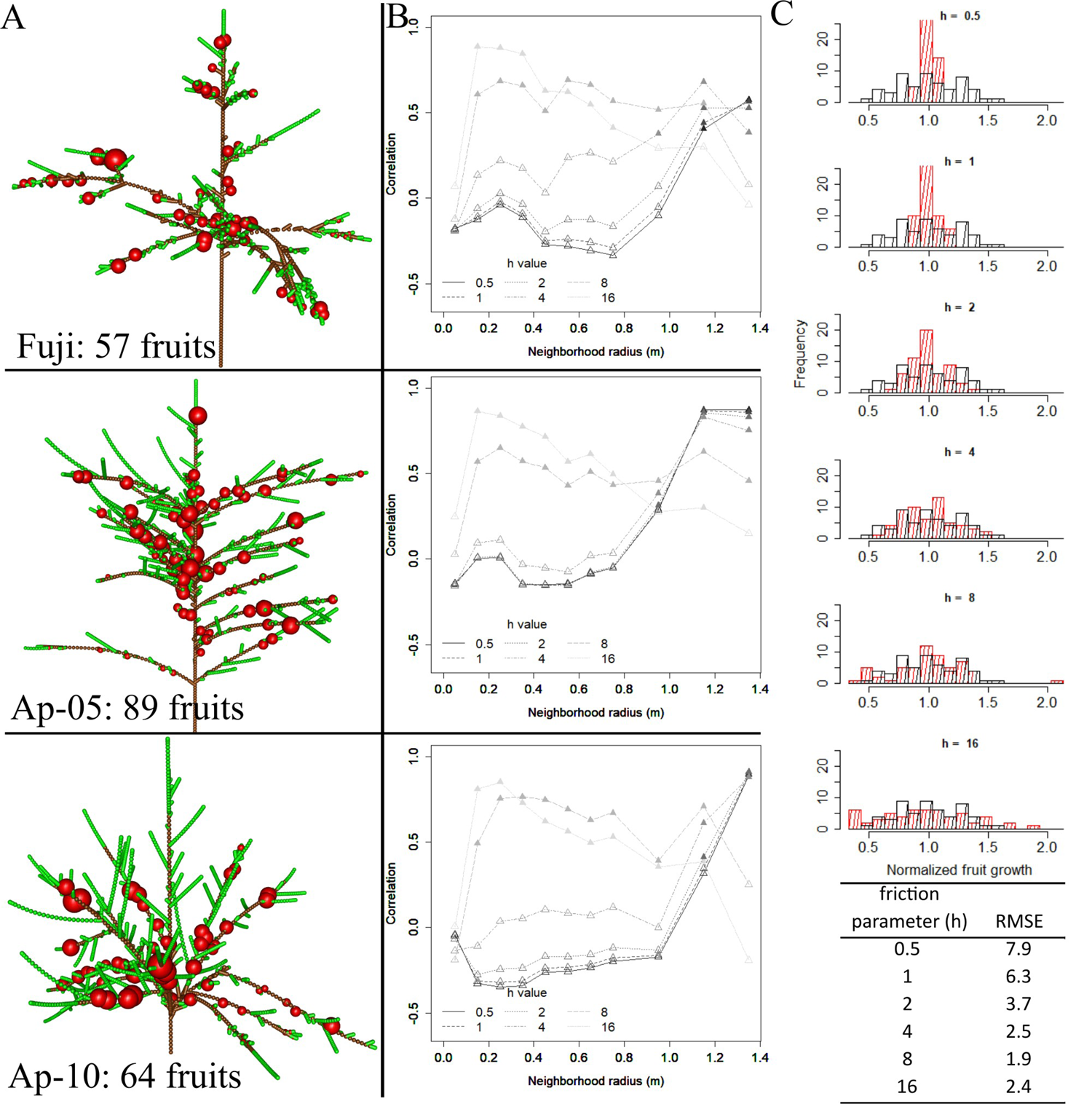
Physiological response of the model run at M scale, in respect to different tree structures and friction parameters (h). A: Three dimensional representation of the tree structures with fruit growth (obtained with h parameter equal to 8). B: correlation between individual fruit growth and the ratio between carbon assimilated and number of fruits evaluated in neighborhoods of increasing radiuses. C: Distibutions of the normalized simulated fruit growth (black, empty bars) superposed with the normalized fruit dry weight measured at harvest (red, coloured bars), in the 4 years old simulated and observed Fuji trees for different friction parameters. The Root Mean Squared Error (RMSE) between distribution counts for different friction parameters is provided. The upper parts of the histograms for h < 2 are cut off in order to allow better visualization of results on the vertical axis.

The normalized fruit dry weight measured at harvest in the four-years old Fuji tree was compared to the simulated fruit in order to evaluate what range of the friction parameter could best reproduce the observed fruit size distribution (Fig.4C). The lowest RMSE values were obtained with h parameters comprised between 4 and 16. High values, however, produced skewed distributions (h = 16) or a variability larger than in the field (h = 8, 16). Conversely, for low h values (0.5, 1, 2) distributions had a consistently lower variability than in the field. Similar variability and low RMSE values were obtained with the friction parameter of four.

### C-allocation at multiple scales

Changing the scale of representation of the tree from M to coarser scales implied a sharp decrease in the number of represented topological components of the tree structures. Trees at GU, TBS, BR1 and FU scales contained respectively about 12.7%, 8.6%, 1.6%, and 1.2% the elements they had at M scale (Table 4). Computation time was found as a third order polynomial function of the number of components in the plant (Fig.5), suggesting that the gain in computation time would increase with the complexity of the plant structure. The reduction in the number of represented plant components, obtained by changing scale from M to FU (down to 0.8%) in the presented simulations, resulted in a gain in computation time of up to four orders of magnitude (down to 0.1%) (Table 4).

**Fig.5:**
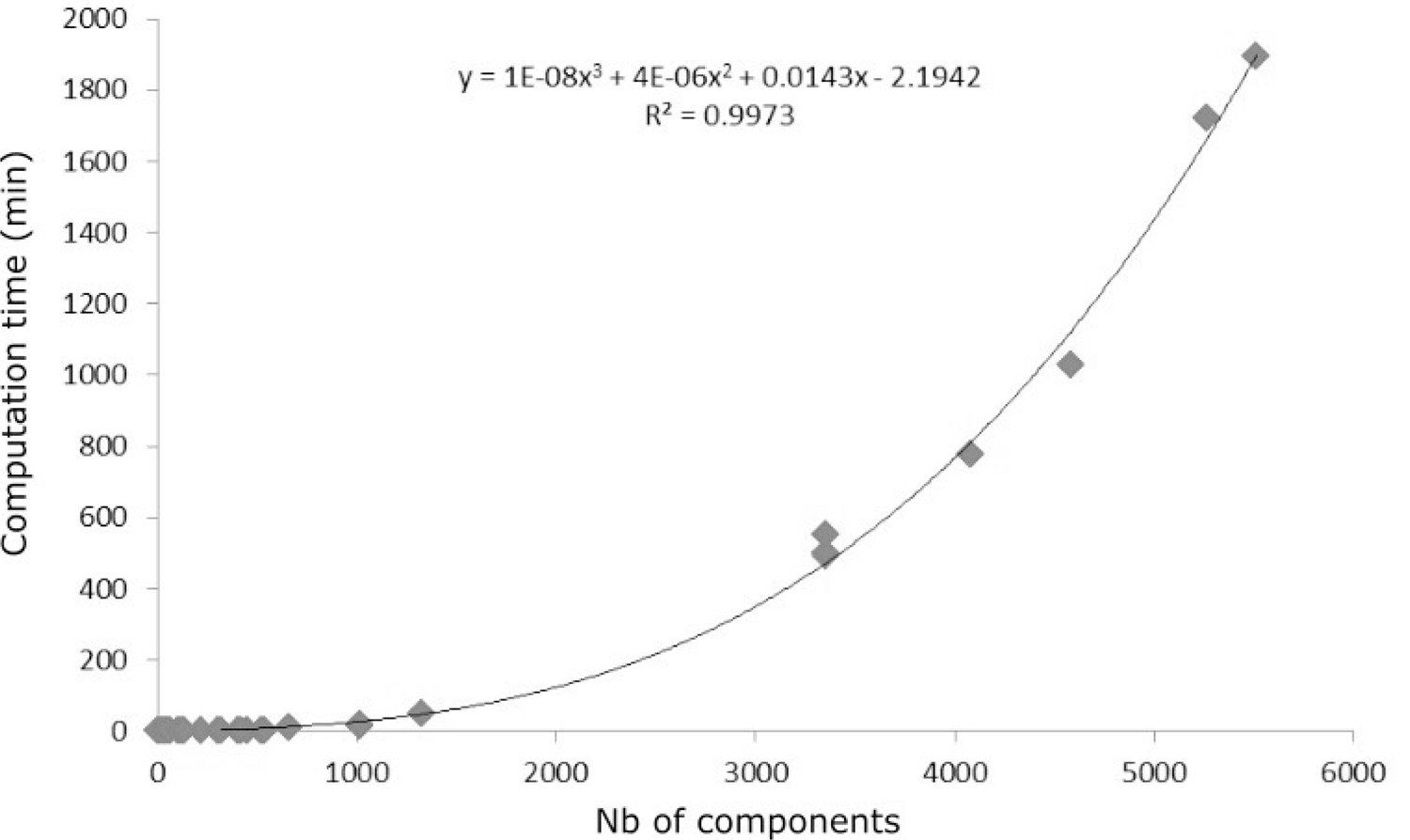
Computation time in respect to the number of represented structure components.

**Table 4:** Number of components represented in tree structures at different scales.

| Tree structure | Scale | Number of components | % | Computation time (min) | % |
| --- | --- | --- | --- | --- | --- |
| Fuji | M | 3355 | 100 | 497.5 | 100 |
|  | GU | 521 | 15.5 | 2.3 | 0.5 |
|  | TBS | 320 | 9.5 | 0.9 | 0.2 |
|  | BR1 | 29 | 0.9 | 0.3 | 0.1 |
|  | FU | 67 | 2 | 0.3 | 0.1 |
| Ap-05 | M | 4579 | 100 | 1026.4 | 100 |
|  | GU | 535 | 11.7 | 2.8 | 0.3 |
|  | TBS | 402 | 8.8 | 2.6 | 0.3 |
|  | BR1 | 115 | 2.5 | 0.7 | 0.1 |
|  | FU | 36 | 0.8 | 0.5 | 0.1 |
| Ap-10 | M | 4083 | 100 | 776.2 | 100 |
|  | GU | 449 | 11 | 2 | 0.3 |
|  | TBS | 302 | 7.4 | 1.2 | 0.2 |
|  | BR1 | 59 | 1.4 | 0.5 | 0.1 |
|  | FU | 36 | 0.9 | 0.5 | 0.1 |

Changing topological scale had also a significant impact on fruit growth. In order to ease the interpretation, only results for the most representative friction parameter values (2 <= h <= 8) are presented in further figures and discussed. Fruit growth at relatively fine scales (GU, TBS) was more correlated to results at M scale than at coarser scales (BR1, FU) (Fig.6). Most often, the higher the friction parameter, the lower the correlation between predictions at any coarse scale and at M scale. However, in some cases (e.g. h = 2 in the the Ap-05 tree structure), the combination of friction parameter and scale resulted in fruit growth predictions systematically different (lower) in respect to the M scale.

As expected, multiple fruits belonging to the same coarse scale component had the same growth (size of spheres on Fig.3). This was due to the fact that the carbon allocated to a coarse scale component is proportionally divided according to individual metamer carbon demand, and all fruits at the beginning of the simulation had identical weight. As a consequence, the higher the resolution in representing the plant structure (Metamer > Growth Unit > TBS > 1^st^ order Branching ≈ Fruiting Unit), the higher the simulated fruit growth variability (Fig.6). Globally, the lower the friction parameter the lower the range of fruit growth variability, in all the tree structures and at all topological scales (Fig.4, Fig.6). The combination of the last two observations is that high friction parameters resulted in relatively wide range of fruit growth at all scales, but with lower variability when moving from a fine to a coarse scale.

**Fig.6:**
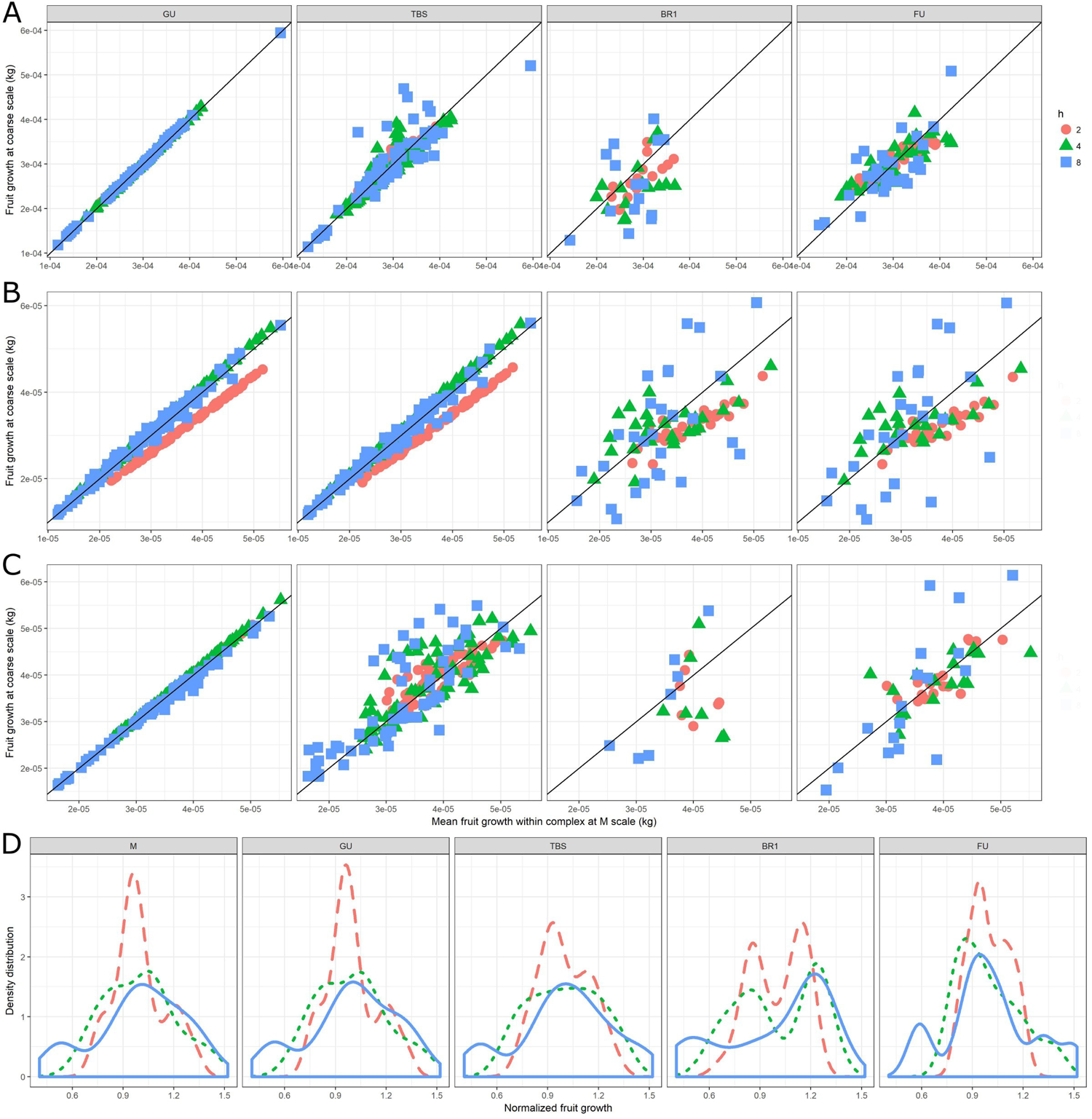
Mean daily fruit dry weights increment and its distribution (in Fuji) depending on the topological scales (from left to right), tree structure (from top to bottom) and friction parameters h (symbols). Correlation between carbon allocated to fruits per day at the selected coarse scale (y axes) vs carbon allocated to fruits per day at M scale and averaged for all metamers belonging to a same component at coarser scale (x axis) in A: Fuji, B: Ap-05, C: Ap-10. D: Fruit growth distribution in the Fuji tree structure (dashed for h=2, dotted for h=4, solid for h=8).

#### Simplification of the tree structures and computation efficiency

The reduction in computation time associated with the use of coarser scales (Table 4) corresponded to an increased discrepancy (in terms of the Coefficient of Variation of the Root Mean Squared Error: CV RMSE) in the results obtained between M and other scales (Fig.7B). Error varied from values next to zero at low friction parameters at GU scale, to values up to thirty-five percent for higher frictions and BR1 or GU scales. This general behavior, however, had a few exception, with a low friction parameter value (2) providing higher discrepancies than higher values at GU and TBS scale in the Ap-05 tree structure.

## DISCUSSION

### Multi-scale coherence, impact on predictions and computation time

To our knowledge, MuSCA is the first C-allocation model able to simulate C-allocation at multiple topological scales within a plant representation. Simulation results revealed that the model was able to produce results highly correlated with the M scale, especially when running at GU scale (Fig.6).

The scale of representation had significant effects on the predicted C-allocation. As a rule of thumb, the deviation between predictions obtained at M and other scales increased when lowering the spatial resolution and for increasing friction parameters (Fig.6, Fig.7). For instance, when running at FU scale, differences in mean fruit dry weight (in terms of coefficient of variation of the RMSE) went up to 60% in respect to the M scale (used in the PEACH model, (Allen *et al.* 2005) (Fig.7B).

**Fig.7:**
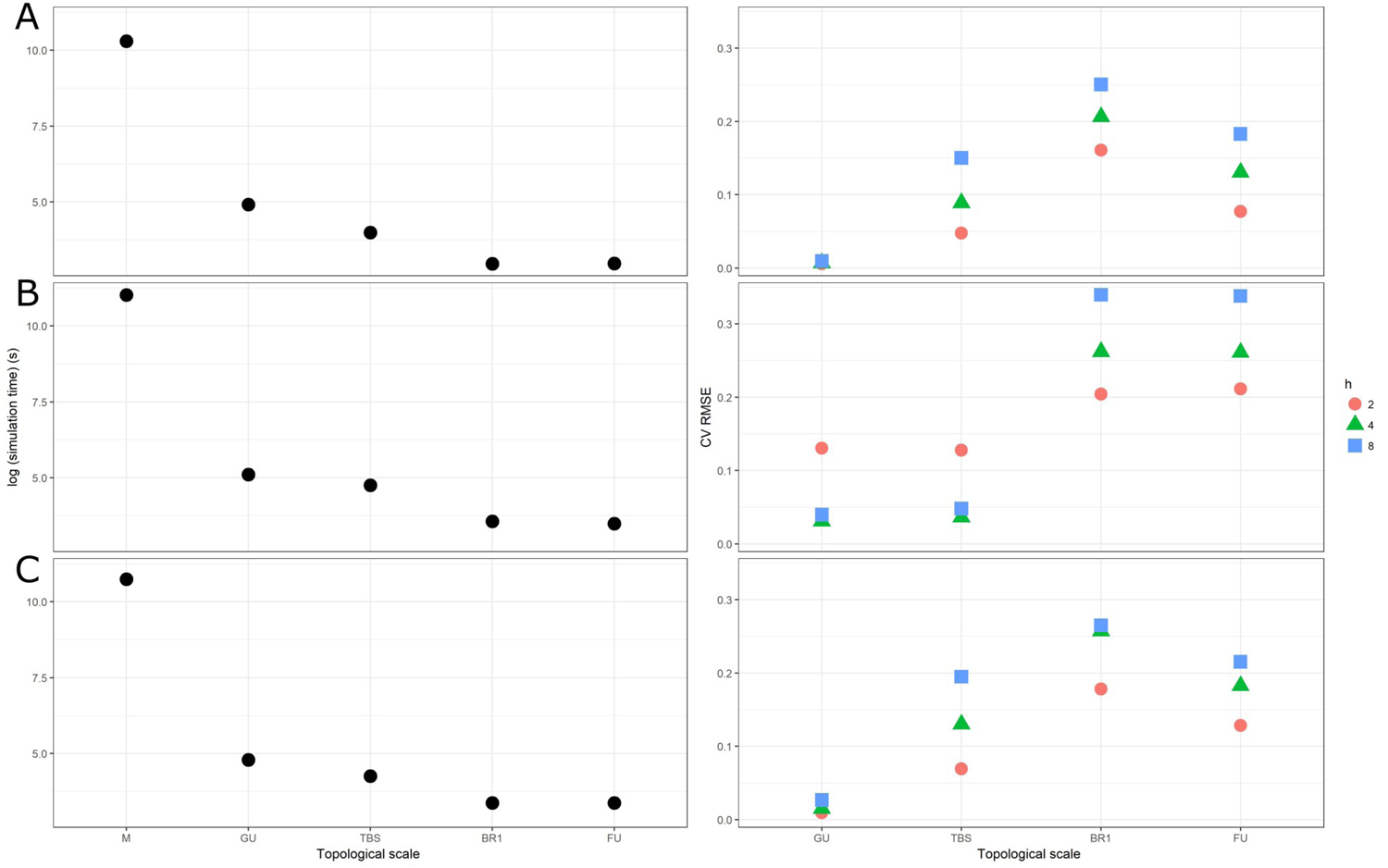
Trade-off between computation time and prediction error. The logarithm of the computation time and the Coefficient of Variation of the Root Mean Squared Error (CV RMSE) between fruit growth at different topological scales in respect to the M scale, on three tree structures and for different friction parameters (h). A: Fuji, B: AP-05, C: AP-10.

Differences in C allocated to fruits at M and other scales are due to two factors. First, while the top and basal coordinates of a coarse scale component are inherited from the finer scale components present at its extremities, the coordinates of its barycentre are not. Indeed these are computed as the mean of the coordinates of its components, weighted by their individual lengths (Calculation of distances). When distances and C-flows are calculated, this generates deviations in respect to the M scale. Second, after allocation, the C received by the different plant parts belonging to a component is not anymore influenced by their individual positions. As such, the model represents the effects of distances among the components at the coarse scale selected for the simulation, but not among its constituting elements at the finer scale. This results in fruits contained in a coarser scale component (of identical initial mass) to grow equally (size of the spheres in Fig.3, e.g. on FU).

The systematic deviations between M and other scales were affected also by the specific plant structure and friction parameter used (Fig.6). The occurrence of systematic deviations is likely related to the non-linearity of the simulated process and its relation to the different discretization of the plant. In trees, C-supplies and C-sinks annual organs (leaves and fruits) alternate with moderate C-sink woody organs creating very contrasted patches in terms of C-supplies and C-sinks (Fig.4). When the same plant is discretized at different scales, the distances between patches of sources and sinks and their constituting elements are modified. By changing discretization, the set of distances to which the non-linear function (eq.1) is applied to compute C flows completely changes, thus implying occasionally sharp differences in C-allocation. Changes are thus related to the geometry of the plant structure as well as to the specific friction parameter, as they determine respectively the spatial domain and the impact of the C-allocation rule. In this regard, further research is needed to explore the impact of commonly applied equations for C-allocation (e.g. eq.1, equations in (Balandier *et al.* 2000; Lescourret *et al.* 2011) on different branching systems in relation to their discretization.

Changing topological scale had a strong impact also on the computation time. This is an important limiting factor for the simulation of carbon allocation in complex tree structures, so that simulations of tree growth at high spatial detail is generally limited to relatively young plants (Balandier *et al.* 2000; Pallas *et al.* 2016). Computation time in MuSCA showed to well fit a third order polynomial function of the number of represented plant components (Fig.5), confirming that minimizing plant complexity can often be a prerequisite to simulate plant growth. In this study, the GU scale was able to reproduce values and fruit growth distributions almost identical to M, while saving computation time (Fig.6, Fig.7). The optimal scale of representation might change with tree size. Especially at early growth stages, relatively small variations in C-dynamics might have important consequences for the tree structure development at later stages. In this regard, we make the hypothesis that intermediate scales comprised between M and GU might further decrease the deviations in respect to M. We further suggest that a mixed use of high and relatively low topological resolutions, respectively for young and older trees, might be a good solution to maximize prediction accuracy while allowing for the simulation of mature trees.

### Testing physiological assumptions

The formalization used in MuSCA to calculate C-allocation (eq.1) (Balandier *et al.* 2000) was able to represent the impact of C-availability and competition among sinks, such as the fruits (Fig.4) on fruit growth variability. The selected formulation presents an advantage, with respect to simpler ones (such as in e.g. QualiTree, (Lescourret *et al.* 2011): its normalization term prevents cases in which the C leaving a plant component is lower or higher than the C-supply available in the source.

Growth at compartment level was in the range of field observations (Fig.4). Relative growth rates obtained at M scale (Table 3) were comprised between normal growth observed in the field and maximum potential growth data used for calibration of sink demands, but were closer to the latter (Reyes *et al.* 2016) between 5.3*10^−3^ and 3.1 *10^−2^ mg/g for the old wood, between 0.4 and 1.2 mg/g for shoots and between 1.9 and 2.8 mg/g for fruits).

In MuSCA, the C-assimilates transportation through the tree structure depends on a friction parameter (*h*, eq. 1). The comparison between the simulated and harvested normalized fruit growth distribution thus suggests that h values suitable for representing a four years old apple Fuji tree should be comprised in the range between two and eight (Fig.4C). This suggests that neither a common assimilate pool (Heuvelink 1995) of carbon nor a shoot or branch autonomy (Sprugel *et al.* 1991) was adequate for accounting for the physiology of assimilate partitioning within a fruit tree as observed in previous experimental studies (Walcroft *et al.* 2004; Volpe *et al.* 2008). Moreover, the existence of fluxes from parts of trees with high carbon supply to parts with a low one have been already described for modeling the within-tree variation in fruit size in peach and apple trees (Lescourret et al. 2011; Pallas et al., 2016). Nevertheless, it has also been observed that the level of shoot or branch autonomy can vary depending on the phenological date with a higher branch autonomy during summer period compared to winter (Lacointe *et al.* 2004). Such kind of behavior could be easily simulated in MuSCA, by dynamically turning the friction parameter values.

A major role of virtual plant growth experiments is to test hypothesis related to the functioning of real plant growth. Indeed, despite the Munch theory was formalized almost ninety years ago, the mechanisms regulating carbon allocation in plants are still the focus of an intense research debate(De Schepper *et al.* 2013). The results presented in this study revealed the impact of the scale of representation on the simulated C-allocation. In this regard, we suggest that the possibility to compare results produced at different scales could help isolating the impact of the scale of representation from the other formalisms (see Fig.6).

### Advantages of flexible C-allocation at multiple-scales

By making a dynamic use of MTG (Godin and Caraglio 1998) the presented model is able to modify the type of “individual entity” represented in a plant structure on the fly. This is possible also thanks to formalisms equally applicable at all scales (Fig.2, eq. 3, 4) that inherit, from fine to coarse scales, the spatial (coordinates of scale boundaries) and extensive (C-demands and supplies, Up-and down-scaling) properties necessary for the calculation of C-flows.

The flexibility related to the multi-scale features of the model presents several opportunities. While the topological scale typically represented in a plant model mirrors the interests of a particular group of model users, the possibility to change scale of representation makes the presented model of practical interest for multiple objectives. The model can be used to assess the implications of using specific scales. As showed in this study, while the use of BR1 or FU scales led to consistent discrepancies in respect to M (up to 35%, Fig.7), at GU the error generally remained much lower (5%). In all cases the reduction in computation time were higher than ninety-nine percent, with minor improvements between GU (99.5%) and BR1 or FU(99.9%) suggesting that, in most cases and for trees of this size, the GU scale would be a robust alternative to M, while allowing much faster simulations.

Comparisons of results with other C-allocation models present in the literature can be facilitated by the possibility to adapt the scale of representation to the one of the other models. By changing the topological scale, the individual fruit growth evaluated at the FU scale can be compared with results obtained by QualiTree (Lescourret *et al.* 2011).

In the presented application, MuSCA was applied on some input MTGs issued by the MAppleT architectural model (Costes *et al.* 2008). However, MTGs coming from other sources, such as several models present in the OpenAlea environment, could also be used, given their compliance with certain prerequisites (Supplementary Information: Inputs of the MuSCA model). If necessary, the detail in the description of the plant could be reduced since, technically, a representation at metamer resolution is not strictly necessary for the functioning of the model. Indeed, in order to run simulations, the plant description should simply allow for the identification of the organ types (via the presence of leaves and fruits), their geometrical sizes (length and radiuses) and their topological connections (succession, branching). This makes the model suitable also for applications on tree structures acquired in the field by methods such as the Terrestrial Laser Scanner (TLS), following topological reconstruction of the plant (Raumonen *et al.* 2013; Boudon *et al.* 2014; Reyes 2016). In addition, because of the flexible representation of the plant structure based on MTGs, field data collected at various topological scales might still be suitable for model testing. Indeed, by using the up- and down-scaling functions, on the one hand results simulated at any spatial scale can be easily brought to the spatial scale at which the data for validation were available.

### Model limitations and further developments

Despite the sound use of a normalization term in the C-allocation rules used in MuSCA, the model still neglects some important physiological implication of the distribution of sinks along the plant topology. In particular, the influence of individual sinks present in the path between sources and sinks on the C-flow is not fully described, as in models based on an electric analogy and the L-system formalisms (Allen *et al.* 2005; Cieslak *et al.* 2011). Indeed, two identical C-sinks (D1, D2) present at the same distance from a source (S) will receive form this source an identical amount of C, no matter if along the path between S and D1 there are stronger or more abundant C-sinks than along the path between S and D2. Nevertheless, simpler models as our has the advantage to require only one parameter to calibrate compared to these previous approaches displaying a larger number of parameters whose calibration could be tedious.

The MuSCA model could be further refined by using the MTG properties to increase the computational efficiency without losing prediction accuracy. For instance, when running at coarse scale, the direction of origin of C-assimilates might be taken into account in order to account for distance also within the boundaries of the coarse scale component.

Despite the presented model was able to represent the effect of competition for C-supplies among C-sinks on various plant structures, some physiological processes essential for the representation of C-dynamics are still not represented. In particular, since the dark respiration alone might account for a loss of about 27% of the CO_2_ during the growing season (Wibbe *et al.* 1993), missing to represent this process certainly leads to important underestimations of the strength of the C-sinks, and thus an underestimation of the competition for C-assimilates. This could explain the reason for not finding competition for C-assimilates in our first simulations, prior to doubling all tree C-demands (Simulation of fruit growth). In addition, the creation of new internodes needs to be included in order to allow for the simulation of plant growth in periods when the plant geometry is not-fixed (shoot elongation period). Further, a more detailed description of the root would be the starting point to investigate water and nutrient limitations at the soil interface.

Regarding the use of MuSCA in the larger context, its genericity (Overview) can ease its adaptation to different species. In addition, the modular implementation of MuSCA in the OpenAlea environment facilitates the integration with other, previously developed, models, as it was the case for the connection with the MAppleT (Costes *et al.* 2008) and the RATP models (Da Silva, Han, and Costes 2014).

## CONCLUSIONS

In this study we presented MuSCA, to our knowledge the first C-allocation model able to simulate C-allocation at multiple topological scales of representation of the plant. The presented model provides topologically-based methods to re-interpret/simplify the topological spatial scale at which the process of carbon allocation is simulated. The model revealed a major impact of the topological scale used to discretize C-sources and sinks on the predicted C-allocation, even when other C-allocation rules (equation for C-allocation and friction parameter) were kept constant. The model can be used to identify what degree of model simplification is acceptable for the representation of plant structures. In other terms it can be used to run simulations, also on large plants, while knowing the trades-off in terms of computation time and prediction accuracy. In addition, the flexible representation of the plant topology permits to more easily meet the needs of different user types, while using the same model. For instance, a relatively coarse scale (e.g. branch) could be more suited for a farmer interested in fruit thinning, than a fine one (e.g. metamer) that could be preferred by a modeler interested in investigating the local drivers of individual fruit size variability. Finally, because of its modular implementation and genericity, minor interventions on a few individual modules are sufficient in order to modify the carbon allocation equation, useful to test other formalisms, or the species specific parameters.

## ACKNOWLEDGEMENTS

We acknowledge Jerome Ngao and Marc Saudreau from UMR PIAF, INRA at Clermont-Ferrand respectively for their help in linking RATP to MAppleT and in linking MTG and RATP, in the OpenAlea platform. F. Boudon from UMR AGAP, Cirad at Montpellier, for some tips on Python coding. The authors thank also Roberto Zampedri and Mr Mauro Cavagna for field assistance and Maddalena Campi for useful discussions.

## Supplementary Information: Inputs of the MuSCA model

Supplementary Information: Inputs of the MuSCA model.
| Variable | Description | Unit |
| --- | --- | --- |
| *Multi-scale Tree Graph description of a plant (.mtg)* |  |  |
|  | Coordinates of the centre of the top of a plant |  |
| XX, YY, ZZ | component | m |
| radius | tree component apical radius | m |
| leaf_area | leaf area surface | m<sup>2</sup> |
| fruit | fresh fruit biomass | Kg |
| observation | organ type: trunk, other types | - |
| Topological |  | - |
| connector | ">" for succession; "+" for branching |  |
| Scale |  |  |
| boundary | "/" for boundary across scales |  |
| *Meteorological file for the RATP model (.mto)* |  |  |
| day | day of the year | day |
| hour | index of the current semi-hour from midnight | 30 min |
| PARglob | global incident Photosynthetically Active Radiation | W m<sup>-2</sup> |
| PARdif | diffused Photosynthetically Active Radiation | W m<sup>-2</sup> |
| NIRglob | global incident Near Infrared Radiation | W m<sup>-2</sup> |
| NIRdif | diffused Near Infrared Radiation | W m<sup>-2</sup> |
| Ratmos | Total incoming radiation | W m<sup>-2</sup> |
| Tsol | mean soil temperature | °C |
| Tair | mean air temperature | °C |
| Eair | H2O partial pressure | Pa |
| CO2air | CO2 partial pressure | Pa |
| Wind | wind speed above tree | m s<sup>-1</sup> |

